# Preservation of cellular nano-architecture by the process of chemical fixation for nanopathology

**DOI:** 10.1101/371286

**Authors:** Xiang Zhou, Luay Almassalha, Yue Li, Adam Eshein, Lusik Cherkezyan, Parvathi Viswanathan, Hariharan Subramanian, Igal Szleifer, Vadim Backman

## Abstract

Transformation in chromatin organization is one of the most universal markers of carcinogenesis. Microscale chromatin alterations have been a staple of histopathological diagnosis of neoplasia, and nanoscale alterations have emerged as a promising marker for cancer prognostication and the detection of predysplastic changes. While numerous methods have been developed to detect these alterations, most methods for sample preparation remain largely validated via conventional microscopy, and have not been examined with nanoscale sensitive imaging techniques. For these nanoscale sensitive techniques to become standard of care screening tools, new histological protocols must be developed that preserve nanoscale information. Partial Wave Spectroscopic (PWS) microscopy has recently emerged as a novel imaging technique sensitive to length scales ranging between 20 and 200 nanometers. As a label-free, high-throughput, and non-invasive imaging technique, PWS microscopy offers many advantages for risk stratification of early cancer, and is an ideal tool to quantify structural information during sample preparation. Therefore, in this work we applied PWS microscopy to systematically evaluate the effects of cytological preparation on the nanoscales changes of chromatin using two cell line models: Hela cells differentially treated with daunorubicin and TP53 differentially mutated ovarian carcinoma cells. Notably, we show that existing cytological preparation can be modified in order to maintain clinically relevant nanoscopic differences, paving the way for the emerging field of nanopathology.

## Introduction

Over the past few decades, despite a tremendous amount of research into discovering new molecular targets and improving precision therapies, cancer remains a leading cause of death worldwide. For almost all types of cancer, treatment effectiveness is directly associated with the stage of detection [1]. Although for low-prevalence malignancies therapeutics remains the primary option for the management of the disease, for more prevalent malignancies such as lung, colon, prostate, and ovarian cancers both the health care costs and mortality rates can be greatly reduced via the development of two-tiered screening strategy. Two-tier screening starts with a cost-effective, patient-compliant, ideally non-invasive or only minimally invasive test that can be administered in the primary care setting and has a sufficiently high sensitivity for clinically significant and treatable lesions. Patients risk-stratified based on this first tier test may then undergo a follow up examination using the more definitive second-tier test. A notable example of the two-tier screening is the pap-smear as a pre-screen for colposcopy paradigm for cervical cancer screening, which after its introduction in clinical care in the 1950s has reduced cervical cancer mortality by more than 95% in the screening population. However, the development of two-tier screening for non-cervical malignancies has been challenging. To date, most attempts to develop this two tier screening methodology have focused on identifying specific molecular transformations correlated with tumor development, with genomic and proteomic markers acting as the two major sources investigated as potential biomarkers[2]. While molecular screening is promising, the heterogeneous accumulation of genetic, epigenetic, and proteomic transformations associated with tumorigenesis make the use of individual markers for screening limited across a wide population. On the other hand, at the later stages of tumorigenesis (e.g. dysplasia and malignancy) these divergent molecular alterations are near universally convergent on microscopic structural alterations that can be identified by the well-established cytological examination. Owing to this convergence between molecular and structural alterations, a number of technologies have been developed for the detection of early stage nanoscopic structural alterations[3]. In these approaches, alterations in nanoscopic ultrastructure act as a convergence point between these numerous independent molecular transformations that are detected by the two-tier screening approach.

One of these methods, Partial Wave Spectroscopic (PWS) microscopy was developed as a label-free, non-invasive optical method to measure nanoscopic changes in cells by analyzing the variations in back-scattered light [4]. With the ability to quantitatively measure length-scales above 20nm, PWS microscopy has demonstrated sensitivity to cellular changes in early and field carcinogenesis for a broad range of cancers, including lung [5], colon [6], ovarian [7], prostate [8], and pancreatic cancer [9]. The most pronounced alterations are observed within the cell nucleus and converge on changes in supra-nucleosomal organization of chromatin which was cross-validated by molecular assays and transmission electron microscopy [10, 11]. As with any cytological microscopy techniques, cells must first undergo fixation and processing in order to maintain their structural stability for long term storage and imaging. Since these preparatory steps are known to have detrimental effects to the cellular structure at the micron-scale, improper preparation could result in nanoscopic distortion and loss of sensitivity to underlying ultrastructural transformations that occur during early carcinogenesis.

In previous work, live-cell PWS microscopy has demonstrated success in visualizing and quantifying cellular changes during chemical fixation, showing that nanoscale structural information can be preserved during chemical fixation methods[12]. Here, we further extended its application to track cellular structure changes during all stages of nanocytological preparation. In particular, we test the effects of a number of common cytological preparations on the ultrastructure of cell lines using baseline live cell measurements as controls. This access into nanoscopic information during all preparation stages starting at live cells allows a generalized frame-work to assess the maintenance of ultra-structure during the preparation of biological samples at all stages of cytological prep: fixation, rehydration and staining. Critically, we find that while some information is lost during nanocytological preparation, clinically relevant information can be relatively well preserved and applied for nanocytological applications for both PWS microscopy as well as for single molecule super-resolution nanoscopy. Thus, the process presented here can be utilized for systematic validation of methods to enable detection of nanoscale structural alterations in human disease to verify that the measured structures represent those underlying pathological process found in live cells.

## Material and methods

### Cell Line Models

Two models were used for the analysis of cytological preparation on cellular ultrastructure: Hela cells differentially treated with daunorubicin in order to cause nucleosomal eviction[13] and human ovarian cancer cell model with differential p53 mutation associated with patient outcome (A2780 and A2780.m248). For each model, nanoscopic changes in chromatin folding were detected and quantified by live-cell PWS microscopy prior to nanocytological preparation. Nucleosomal eviction in HeLa cells was induced using 10μM daunorubicin treatment, with nanoscopic changes observed in live cells within 15 minutes. Comparatively, a model of tumor aggressiveness was analyzed using the ovarian A2780 cancer model in comparison with the p53 mutated cell line A2780.m248 (M248). As p53 has been shown to alter gene expression by changing chromatin organization through the post-translational modifications of histones, this mutation to the core binding p53 domain could confer differential ultrastructure that is associated with worse prognostic outcome [14].

### Cell culture

HeLa cells and human ovarian cancer cells A2780 and A2780.m248 (M248) were grown in freshly prepared RPMI-1640 media (Life Technologies) supplemented with 10% (vol/vol) FBS (Sigma-Aldrich) and grown at 37°C and 5% CO_2_. All of the cells in this study were maintained between passage 5 and 30. Microscopy measurements were obtained from cells grown on uncoated size 1 glass coverslips attached to 50mm petri dishes (Cell Vis). Petri dishes were seeded with between 10,000 and 50,000 cells in 2ml of the cell appropriate media at the time of passage. Cells were allowed at least 24 hours to re-adhere and recover from trypsin-induced detachment. Imaging was performed when the surface confluence of the slide was between 50-70% and when cells were in the cell culture media. A reference scattering spectra was obtained from an open surface of the substrate coverslip immersed in media to normalize the intensity of light scattered for each wavelength at each pixel.

### Chemical Fixation

After live cell imaging, cell culture media was removed from the petri-dish and cells were washed with 2mL phosphate buffered saline solution twice. After removal of the washing solution, 2mL of specified fixative solution was added into the dish. Six common fixatives were tested in our study: 1. acetic acid: ethanol = 1:3 (v/v%); 2. Carnoy’s fixative (Ethanol : chloroform : acetic acid = 6:3:1 (v/v%)); 3. FAA fixative (Ethanol : formaldehyde : acetic acid=16:3:1 (v/v%)); 4. 4% formaldehyde in PBS solution (pH~7.4); 5. 2.5% glutaraldehyde and 2% formaldehyde in PBS solution (pH~7.4). 6. 95% ethanol (v/v%). After 15 minutes of fixation at room temperature, the cells were imaged in the fixative solution. A reference spectra of the coverslip immersed in the specified fixatives solution was obtained for normalization.

### Serial rehydration

After the 95% ethanol fixation, cells were sequentially rehydrated with 2mL of 70%, 50%, 25% ethanol solution and DI water for 10 minutes each under room temperature. The same cells were imaged in the specified solution after each step of rehydration. For comparison, cells were directly rehydrated with DI water after 95% ethanol fixation and were imaged in DI water after 30 minutes. After each step of imaging, a reference spectrum of the coverslip immersed in the specified solution was obtained for normalization.

### Air drying

After serial rehydration, DI water was removed from the petri-dish and 100mM trehalose-water solution was added into the petri-dish as a drying solution. Trehalose is hypothesized to prevent cell shrinkage and protect cell membranes by forming high-viscosity glass matrix during evaporation of solvents. It has been reported that cells with intra cellular trehalose has improved tolerance against freezing and desiccation. [15, 16] In our study, since cells were fixed and their membrane integrity was no longer maintained, trehalose will penetrate into cells by diffusion and serve as structural protection molecules during air drying. After a 30-minute treatment, trehalose solution was completely removed from the petri-dish by pipetting. Cells were then air dried at room temperature for 48 hours. In addition, we investigated the effects of air drying speed on chromatin organization as measured by PWS. For humidity-controlled air drying, cells were air dried without trehalose treatment under three humidity conditions (25%, 50% and 75%) at room temperature. After air drying, cells were imaged in 95% ethanol. A reference spectra of the coverslip immersed in the 95% ethanol solution was obtained for normalization.

### Traditional histological staining

After air drying and imaging, cells were stained with Hematoxylin and Cyto-Stain (Thermo Scientific, Richard-Allan Scientific) in the petri-dish. The staining process consisted of washing with DI water for 60 seconds, staining with Hematoxylin II for 25 seconds, washing with DI water for 15 seconds, washing with clarifier for 45 seconds, washing with DI water for 30 seconds, washing with bluing reagent for 25 seconds, washing with DI water for 30 seconds, washing with 95% ethanol for 30 seconds, staining with Cyto-Stain for 25 seconds and final washing with 95% ethanol for 60 seconds. After staining, the same cell populations were imaged again in 95% ethanol. A reference spectra of the coverslip immersed in the 95% ethanol solution was obtained for normalization.

### Immunofluorescence staining

The immunofluorescence staining was performed on HeLa model. The staining process consisted of: 10 minute PFA fixation (4% formaldehyde in PBS solution, Electron Microscopy Sciences), washing with PBS solution, blocking and permeabilization (1% BSA and 0.1% Triton X-100 in PBS) for 20 minutes, washing with PBS solution, incubation with primary antibody (Anti-Histone H3K9 me3, Abcam) in blocking solution at 4°C overnight, washing with blocking solution, incubation with secondary antibody (Goat anti mouse Alexa Fluor 488) in blocking solution for 2 hours and final washing with PBS solution. Hoechst 33342 staining was performed after the immunofluorescent preparation for co-localization. PWS measurements of the same cells were taken in PBS solution after each wash step and a reference spectra of the coverslip immersed in the PBS solution was obtained for normalization. Fluorescent images of the same cells was performed at the end of the preparation.

### Immunofluorescence staining for STORM/PALM nanoscopic imaging

The immunofluorescence staining was performed on human ovarian cancer model. The staining process consisted of 10 minute PFA fixation (4% formaldehyde in PBS solution, Electron Microscopy Sciences), washing with PBS solution, reduction in 0.5% sodium borohydride in PBS to reduce auto-fluorescence from the background, blocking and permeabilization (1% BSA and 0.1% Triton X-100 in PBS) for 20 minutes, washing with PBS solution, incubation with primary antibody (Anti-RNA polymerase II, Abcam) in blocking solution at 4°C overnight, washing with blocking solution, incubation with secondary antibody (Alexa Fluor 546) in blocking solution for 2 hours and final washing with PBS solution. PWS measurements of the same cells were taken in PBS solution after each wash step and a reference spectra of the coverslip immersed in the PBS solution was obtained for normalization. STORM imaging was performed at the end of the preparation.

## Results

### PWS imaging of live cell models

At any given intercellular location, refractive index *n* is proportional to the local macromolecular density (proteins, DNA, RNA, etc.) (*ρ*): *n*(***r***) = *n_water_* + *αρ*(***r***) with *α* the refraction increment, which is nearly constant for most kinds of macromolecules[17]. Spatial variations in refractive index result in light scattering, and thus variations in cellular (e.g. chromatin) density can be assessed through the analysis of light scattering properties of a cell. Live-cell PWS microscopy measures the optical interference signal of the backscattering light that is produced by spatial variations of refractive index [18-20]. By quantifying the standard deviation (Σ) of the spectra acquired by PWS microscopy, PWS is able to obtain subdiffractional information from cellular structures, in particular the internal structures of nucleus. Σ is a measure of the heterogeneity of macromolecular density *ρ*(***r***) and is proportional to the fractal dimension (packing scaling) D of the chromatin density variations within a given coherence volume: Σ ~ *D* − *D*_0,_ where *D*_0_ ~ 1.5 is close to the minimal fractal dimension (5/3) that a polymer can attain in 3D space [21, 22]. *D* characterizes the type of scaling between the mass of chromatin contained (*M*) within a sphere or radius *R*: *M* ∝ *R^D^*; *D*<3 for fractal scaling (in which case *D* is referred to as the mass fractal dimension), which is typical for polymeric structures such as chromatin. Length scale sensitivity of PWS depends on the illumination and light collection geometry of the microscope and is typically optimized to sense the chromatin length scales that correspond to the supranucleosomal chromatin structure from the size of chromatin chains to the size of topologically associated domains are the most significantly altered in early carcinogenesis (20-350nm; from ~1kb to 1–10Mbp) [21] [23] [24]. PWS generates an image of a cell such that for each diffraction-limited pixel within the cell nucleus nanoscale heterogeneity of chromatin packing is measured as either Ʃ or *D*.

For clinical diagnostic applications, the use of live cells is problematic and fixed cytology or histology specimens are more often used. To study nano-cytological preparation protocols to preserve the nanoscopic chromatin structure, two cell line models were imaged by live-cell PWS at live state as controls (Fig. 1A, 1B). The cell nuclei in each group were segmented for mean nuclear Σ quantification. The mean nuclear Σ was normalized and rescaled to 100% on average with respect to the appropriate control group (untreated HeLa cells and A2780 cells for all cytological conditions). The mean nuclear Σ of HeLa cells showed a 31% decrease after the 15min 10μM daunorubicin treatment as compared to the control group (P < 0.001). Conversely, the mean nuclear Σ of M248 displayed a 17% increase due to the p53 mutation as compared to the wild-type A2780 (P < 0.001) (Fig. 1C). As these alterations in higher-order chromatin organizations (ΔΣ) are introduced by distinct mechanisms, they can act as molecularly distinct models of nanoscopic changes of chromatin topology to assess the effects of nanocytological preparation.

**Figure 1.**
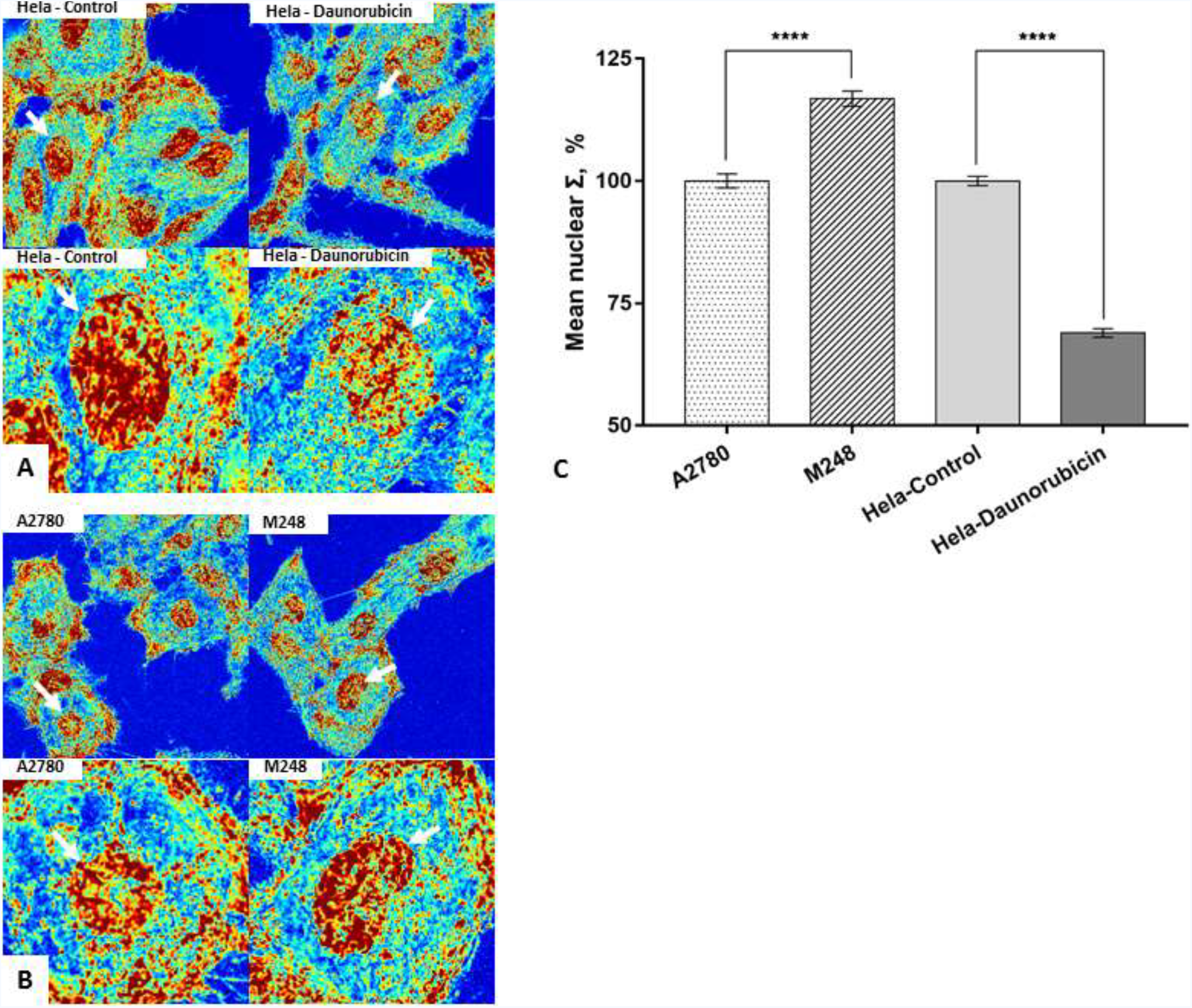
Molecularly distinct structural models of nanoscopic changes to chromatin topology. (A) Representative images of HeLa cells before and 15 min after daunorubicin treatment. Σ was scaled to range between 0.005 and 0.065. (B) Human ovarian cancer cells M248 and A2780. Σ was scaled to range between 0.005 and 0.045. (C) Quantification of mean nuclear Σ change in HeLa model before and 15 min after daunorubicin treatment (HeLa-control=201 cells, HeLa-daunorubicin=200 cells), and in human ovarian cancer model due to p53 mutation (M248=240 cells, A2780=228 cells) with SE bars.

### Effects of different chemical fixation methods on nuclear structure

To determine the effects of chemical fixation methods on our structural models, the same cell populations were imaged before and after fixation, and their mean nuclear Σ was quantified. For 95% ethanol fixation, slight morphological changes were observed after 15 minutes, but the cell nucleus remained clear and detectable in both models. The ΔΣ was quantified to be ~20% for HeLa model (P < 0.001), and ~17% (P < 0.001) for ovarian cancer model (Fig. 2C). Our results showed that the nanoscopic structural differences (ΔΣ) between populations in both models remained significant during 95% ethanol fixation. Although the population difference was preserved after fixation, on the single cell level, we found a weak correlation between nuclear structure before and after fixation (Pearson correlation coefficient (PPC) = ~ 0.4) (Fig. S2). This implies that during fixation different individual nucleus could undergo very different structural transformations, such that the single cell level difference might not be reliably detectable after fixation. In addition, we tested five fixatives that were commonly used in traditional histology studies or immunofluorescent microscopy. As can be seen from Figure 3, the crosslinking fixatives, formaldehyde and glutaraldehyde were able to preserve the ΔΣ, while other fixatives that contain acetic acid resulted in ΔΣ loss after 15 minutes of fixation. Since 95% ethanol is a standard cytological fixative and is stable over longer periods than the other fixatives, the subsequent nanocytology protocols (rehydration, air drying and traditional histological staining) were developed and tested based on 95% ethanol fixation. On the other hand, crosslinking fixation was used for immunofluorescent labeling as the most common approach.

**Figure 2.**
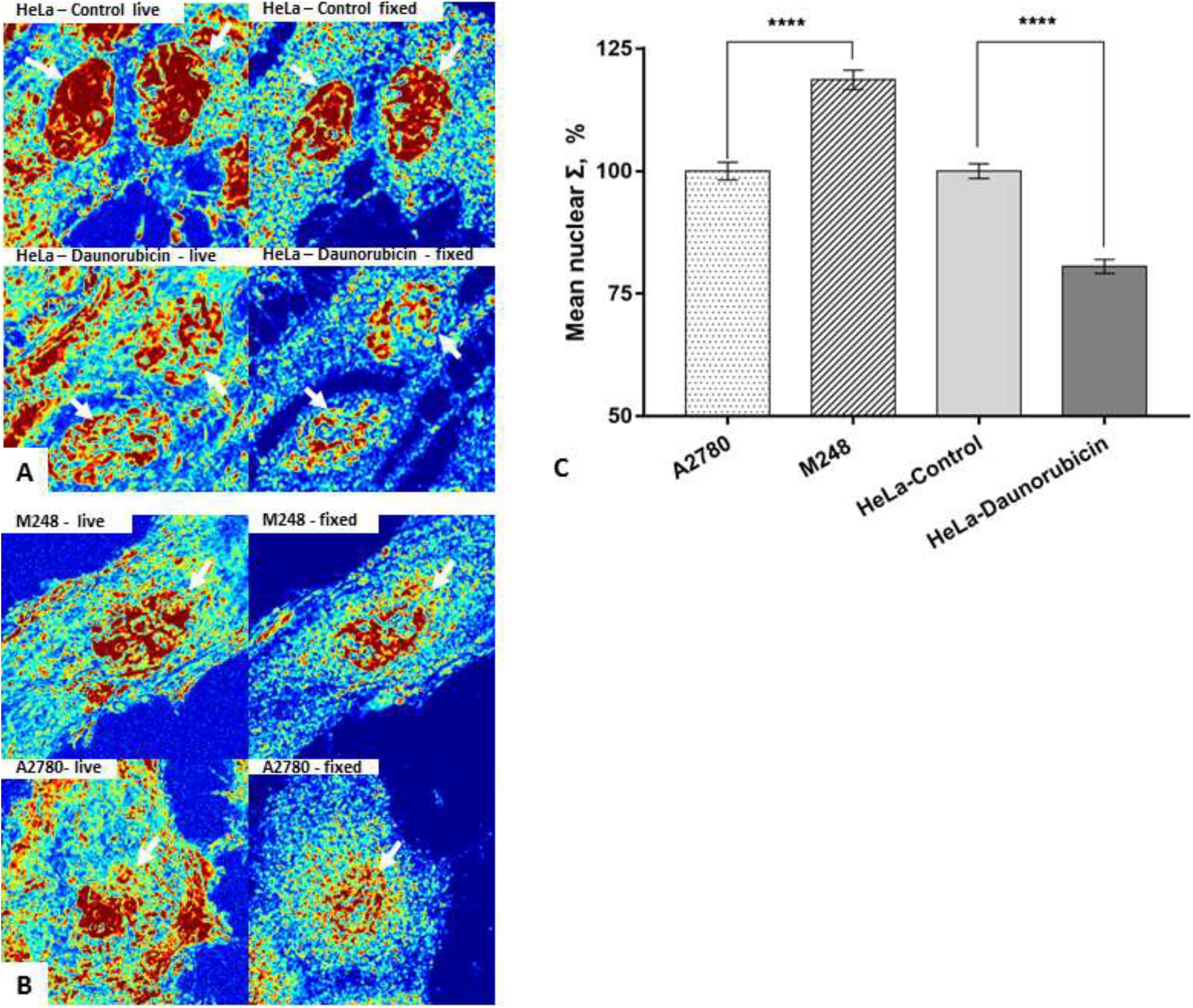
Preservation of ΔΣ by 95% ethanol fixation in two structural cell line models. (A) Control and daunorubicin treated HeLa cells before and after 95% ethanol fixation. (B) Human ovarian cancer cells M248 and A2780 before and after 95% ethanol fixation. Σ is scaled to range between 0.005 and 0.045 for live cells and is scaled to range between 0.02 and 0.14 for fixed cells. (C) Quantification of mean nuclear Σ difference in HeLa cell model (HeLa-control=141 cells, HeLa-daunorubicin=145 cells), and in human ovarian cancer cell model after 95% ethanol fixation (M248=110 cells, A2780=115 cells) with SE bars.

**Figure 3.**
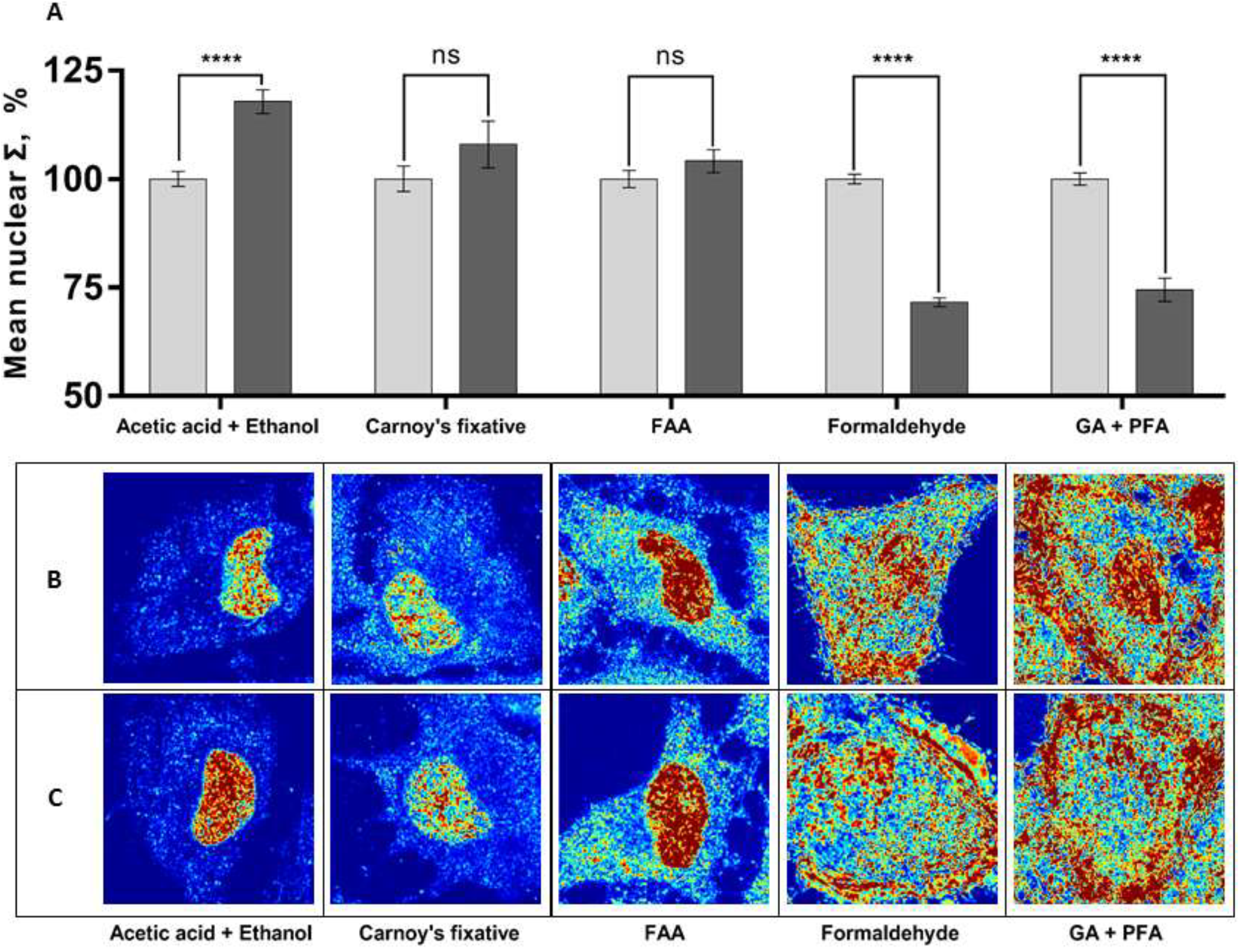
Commonly used fixatives and their effects on ΔΣ in HeLa model. (A) Quantification of mean nuclear Σ ((HeLa-control (grey columns), HeLa-daunorubicin (black columns)) before and after fixation using five different fixatives: 1. acetic acid: ethanol = 1:3 (v/v%), n=~80 cells; 2. Carnoy’s fixative (Ethanol : chloroform : acetic acid = 6:3:1 (v/v%)), n=~30 cells; 3. FAA fixative (Ethanol : formaldehyde : acetic acid=16:3:1 (v/v%)), n =~40 cells; 4. 4% formaldehyde in PBS solution (pH~7.4), n=~90 cells; 5. 2.5% glutaraldehyde and 2% formaldehyde in PBS solution (pH~7.4), n=~60 cells. (B) Representative PWS images of HeLa-control (top row), HeLa-daunorubicin (bottom row) fixed by different fixatives.

### Serial rehydration

Although 95% ethanol preserves the ΔΣ, it could cause cell shrinkage and aggregation especially when cells were freely suspended in solution. To address this problem, we rehydrated the 95% ethanol-fixed cells with gradually decreasing concentrations of ethanol. PWS images of the same cells were acquired at each ethanol concentration condition during the rehydration process. For both structural models, we observed a slight increase in Σ in cytoplasmic regions after serial rehydration, but the cell nucleus remained detectable throughout the whole process (Fig. 4B, 4C, 4E and 4F). By quantifying the mean nuclear Σ in each step of rehydration (Fig. 4A and 4D), we found that the structural differences (ΔΣ) in both models were well preserved. In comparison to this serial rehydration, a direct rehydration from 95% ethanol fixation to DI water was also tested on HeLa model (Fig. 4G and 4H) and resulted in a total loss of measureable differences between the samples. This differential response in structure to the solvation suggests that a rapid shifts in the osmotic pressure by switching solvents could alter the observed differences in the chromatin structure.

**Figure 4.**
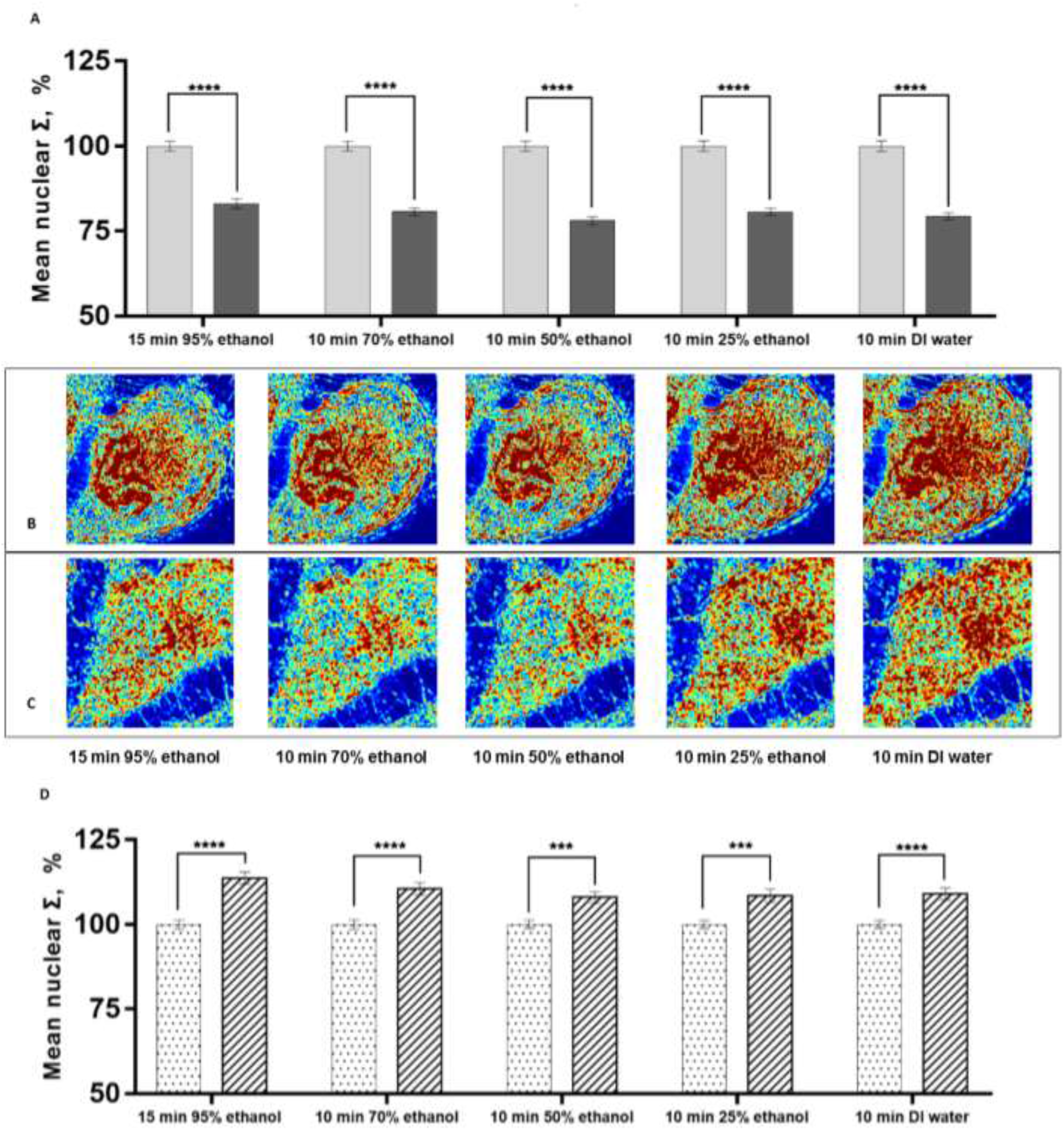
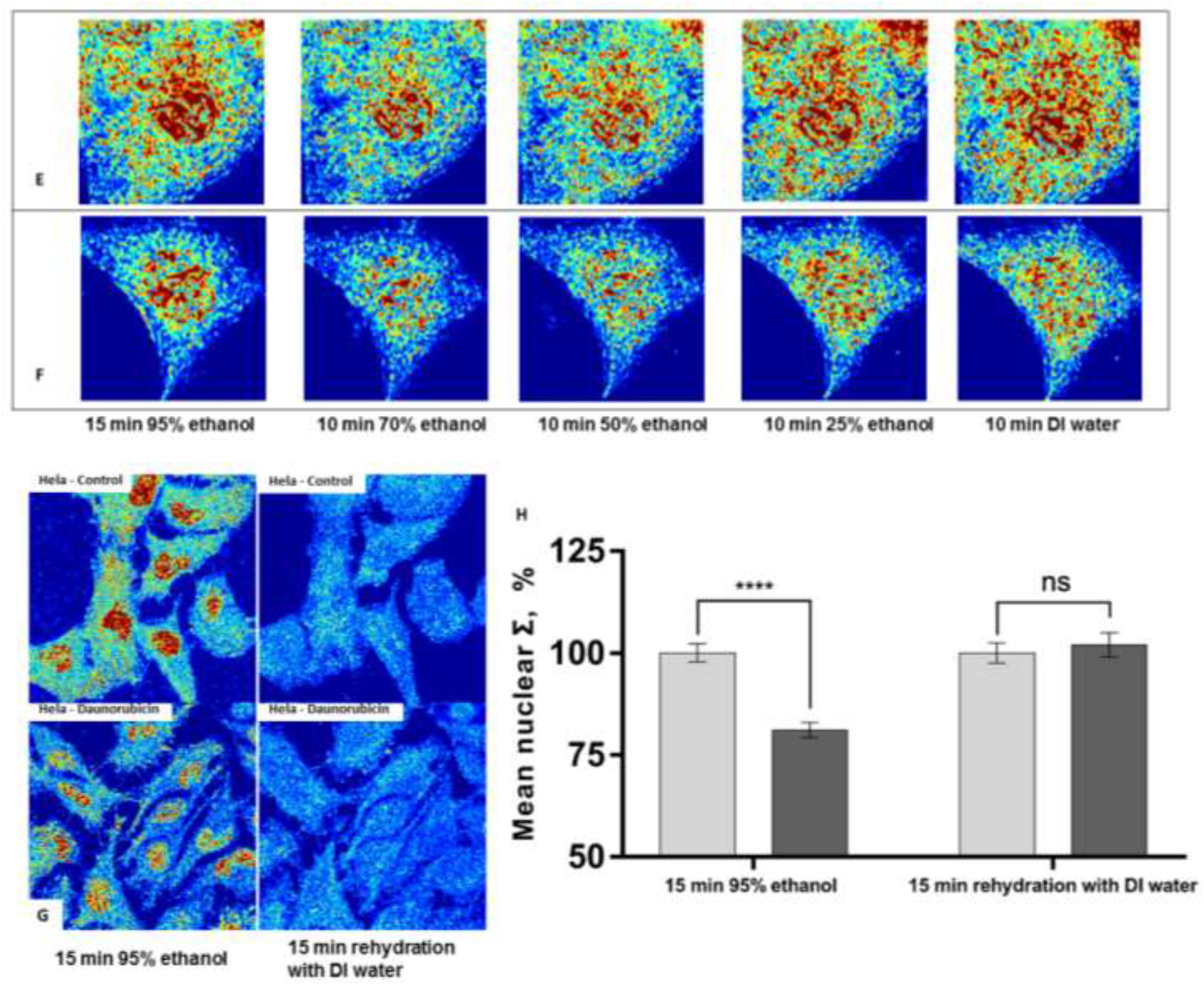
The effects of rehydration on ΔΣ in two structural cell line models. Quantification of Σ in HeLa cell model (A) (HeLa-control (grey columns) =125 cells, HeLa-daunorubicin (black columns) =125 cells), and in human ovarian cancer cell model (D) (M248 (columns with dots) =61 cells, A2780 (columns with stripes) =67 cells) after each step of serial rehydration with SE bars. The ΔΣ was preserved during serial rehydration in both models. Representative PWS images of the same HeLa cells (control (B) and daunorubicin treated (C)), and human ovarian cancer cells (M248 (E) and A2780 (F)) after each step of serial rehydration. Σ is scaled to range between 0.02 and 0.14. (G) Direct rehydration was performed on HeLa model, and the same cells were images by PWS before and after direct hydration with DI water. Σ is scaled to range between 0.02 and 0.14. (H) Quantification of Σ (HeLa-control (grey columns) =42 cells, HeLa-daunorubicin (black columns) = 46 cells) showing the loss of ΔΣ after direct rehydration.

### Preservation of ultrastructure during air drying

Air drying is a common preparatory process in cytology because it allows the adhesion of cells to the substrate and stabilizes cellular structures onto the glass substrate. However, air drying is also known to cause cell volume changes as well as internal structural distortions. To preserve the nanoscopic structural information for PWS imaging after the process of air drying, we treated the rehydrated cells with trehalose solution for 30 minutes before air drying. After 48-hour air drying, the same cell populations were imaged again in 95% ethanol solution (Fig. 4A and 4B), and their mean nuclear Σ was quantified (Fig. 5D). Our results suggest that air drying from trehalose solution maintained the cell morphology and preserved the ΔΣ in both models (~20% for Hela model, P<0.001 and ~12% for ovarian cancer model, P<0.001). The preservation of ΔΣ could be explained by the formation of highly viscous trehalose glass inside the cells which immobilized the macromolecules and stabilized the internal structures during evaporation of water. In comparison, we investigated the effects of direct air drying under different humidity conditions without trehalose treatment. As can be seen in Fig. S5, air drying in low (~25%) and medium (~50%) humidity condition preserved the ΔΣ, but they both resulted in substantial morphological changes and great reduction in ΔΣ. This result indicates that direct air drying which would be detrimental to preservation of cellular ultrastructures.

**Figure 5.**
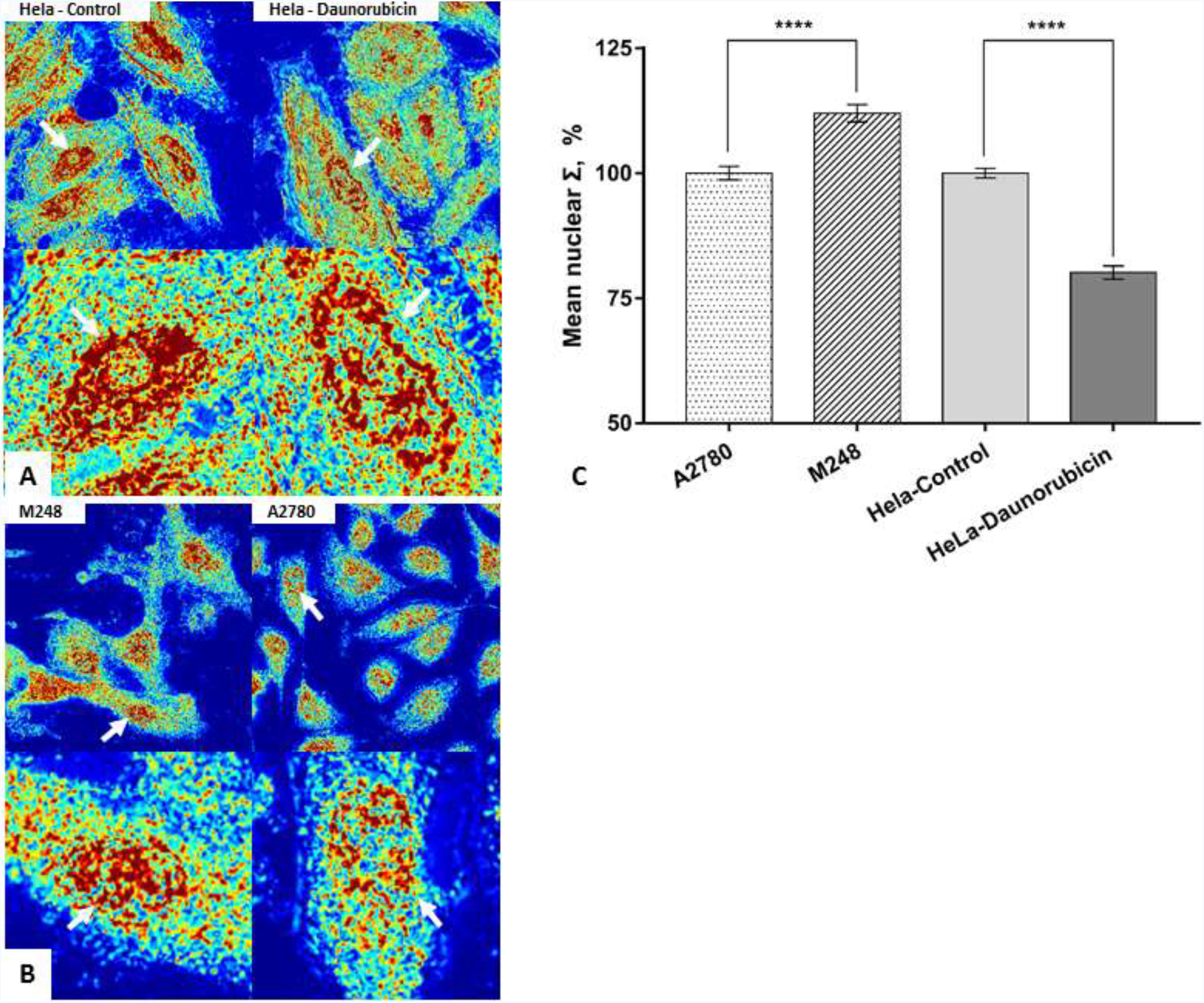
Preservation of ΔΣ after air drying from trehalose solution in two structural cell line models. (A) HeLa model and (B) human ovarian cancer model imaged by PWS after air drying. Σ is scaled to range between 0.02 and 0.25. (C) Quantification of mean nuclear Σ in HeLa cell model (HeLa-control=151 cells, HeLa-daunorubicin=140 cells, p<0.001), and in human ovarian cancer cell model (M248=62 cells, A2780=60 cells, p<0.001) with SE bars.

### Histological staining

After air drying from trehalose solution, we stained the two cell line models with Hematoxylin and Cyto-Stain. In histology, Hematoxylin is a commonly used nuclei stain and Cyto-Stain is mix of dyes used for polychromatic staining. The staining process consists of multiple steps (see M&M) which might alter the chromatin organizations. To verify that the staining process does not result in a loss in diagnostic performance, we imaged the same cell populations with PWS microcopy throughout all stages. As can be seen in Figure. 6, the process of staining results in a clearer demarcation of the nucleus while maintaining diagnostic differences (ΔΣ) were preserved for both models. However, the reduction in the size of ΔΣ in both models also indicate that the staining process could cause structural information loss.

**Figure 6.**
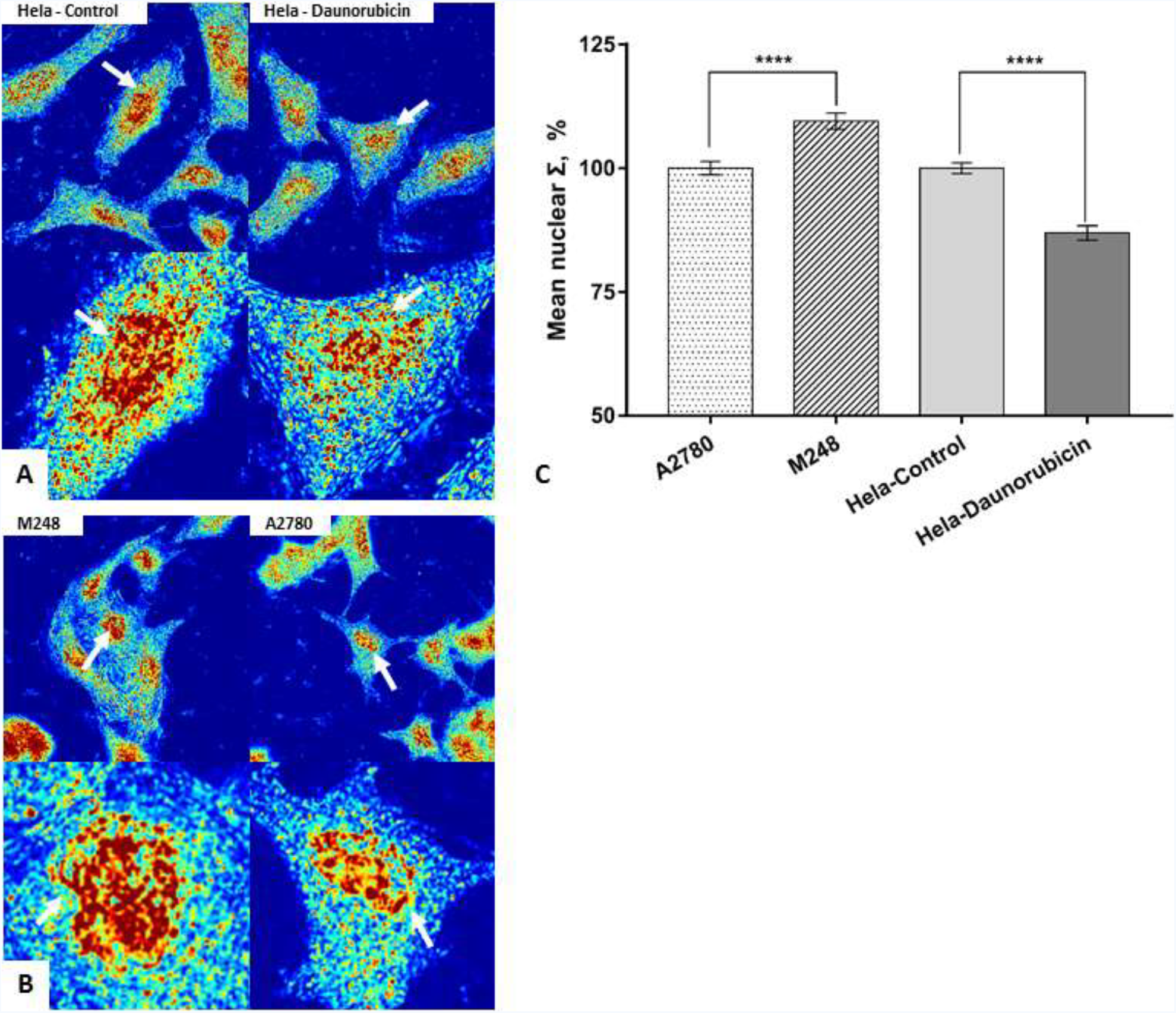
Effects of Hematoxylin and Cyto-Stain on ΔΣ in two structural cell line models. Representative PWS images of HeLa model (A) and human ovarian cancer model (B) after staining. (C) Quantification of mean nuclear Σ in HeLa model (HeLa-control=82 cells, HeLa-daunorubicin=75 cells, p<0.001), and in human ovarian cancer model (M248=73 cells, A2780=76 cells, p<0.001) with SE bars.

### Immunofluorescent labeling

Immunofluorescence (IF) staining is a widely used technique that allows target structures to be visualized and located by light microscopy. In recent years, although many improvements in fluorophores and detection methods have been made, the major procedures of immunofluorescent labeling remained unchanged. These procedures, including chemical fixation, blocking, permeabilization, antibody incubation and multiple steps of washing, are known to cause structural distortions in the cell. Here, we performed PWS imaging at each step of the fluorescent labeling to study the nanoscale changes in chromatin organization. Fluorescent and STORM images were also acquired at the end of the preparation to verify the labeling process (Fig. 7 and 8). Quantitatively, the structural differences (ΔƩ) in both models were preserved during each preparatory step. Cell morphology was maintained. Further, we performed co-localization analysis of PWS and fluorescent images on HeLa model (Fig. 7D). Since the cells were labeled with anti-H3K9me3 antibody which is an indicator of the heterochromatin and denser regions of the nucleus, in theory the mass density distribution of these subdivisions (as indicated by Σ) could be related to their fluorescent labeling density. To test our hypothesis, we first divided the pixels within each nuclei equally into 10 subdivisions based on their fluorescent intensity rankings. And then, we calculated the relative Σ of the subdivisions by normalizing their average Σ to the average Σ of each nuclei. Finally, the relative Σ for each subdivision was averaged across 316 cells. For 316 cells analyzed, the relation between fluorescent intensity and the relative nuclear Σ follows an exponential decay model (R^2^= 0.955):

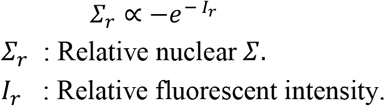

**Figure 7.**
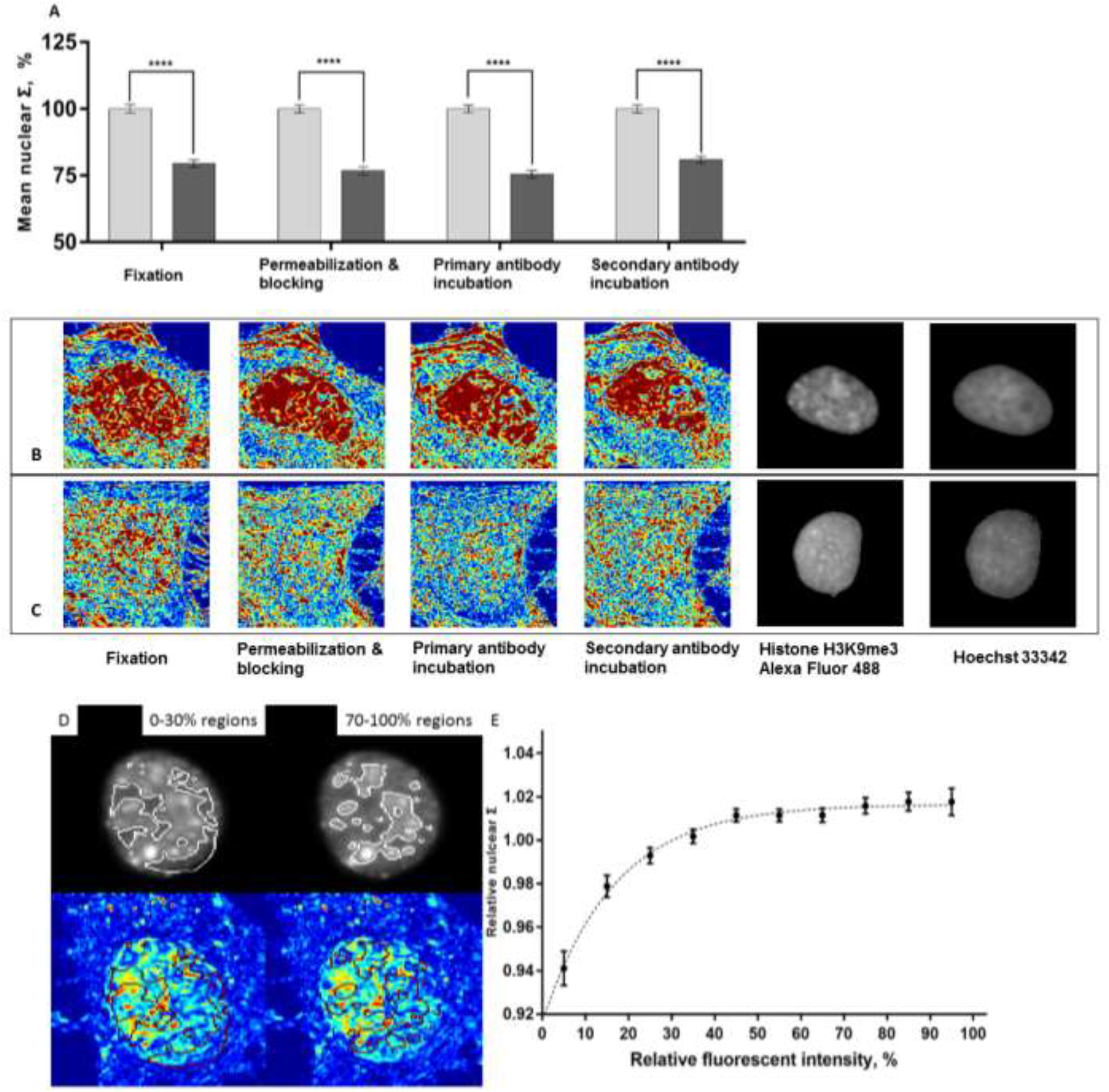
Effects of immunofluorescence staining on ΔΣ. HeLa cells were stained using antibody for H3K9me3 (Alexa Fluor 488). (A) Quantification of Σ in HeLa cell model (HeLa-control = 72 cells, HeLa-daunorubicin=78 cells) at each step of immunofluorescent staining. Representative PWS (left) and fluorescent images (right) of HeLa-control (B) and HeLa-daunorubicin (C) at each step of immunofluorescent staining. (D) Colocalization using PWS and fluorescent microscopy. Each nucleus was segmented into 10 subdivisions based on their fluorescent intensity rankings. Regions with 0-30% and 70-100% fluorescent intensity rankings were shown on both fluorescent and PWS images. (E) Relationship between relative average Σ and fluorescent intensity, with standard error. For each nucleus, we calculated the relative Σ of each fluorescent subdivision by normalizing its Σ to the average Σ of the whole nucleus. The graph shows the averaged relative Σ for each fluorescent subdivision across 316 nucleus.

**Figure 8.**
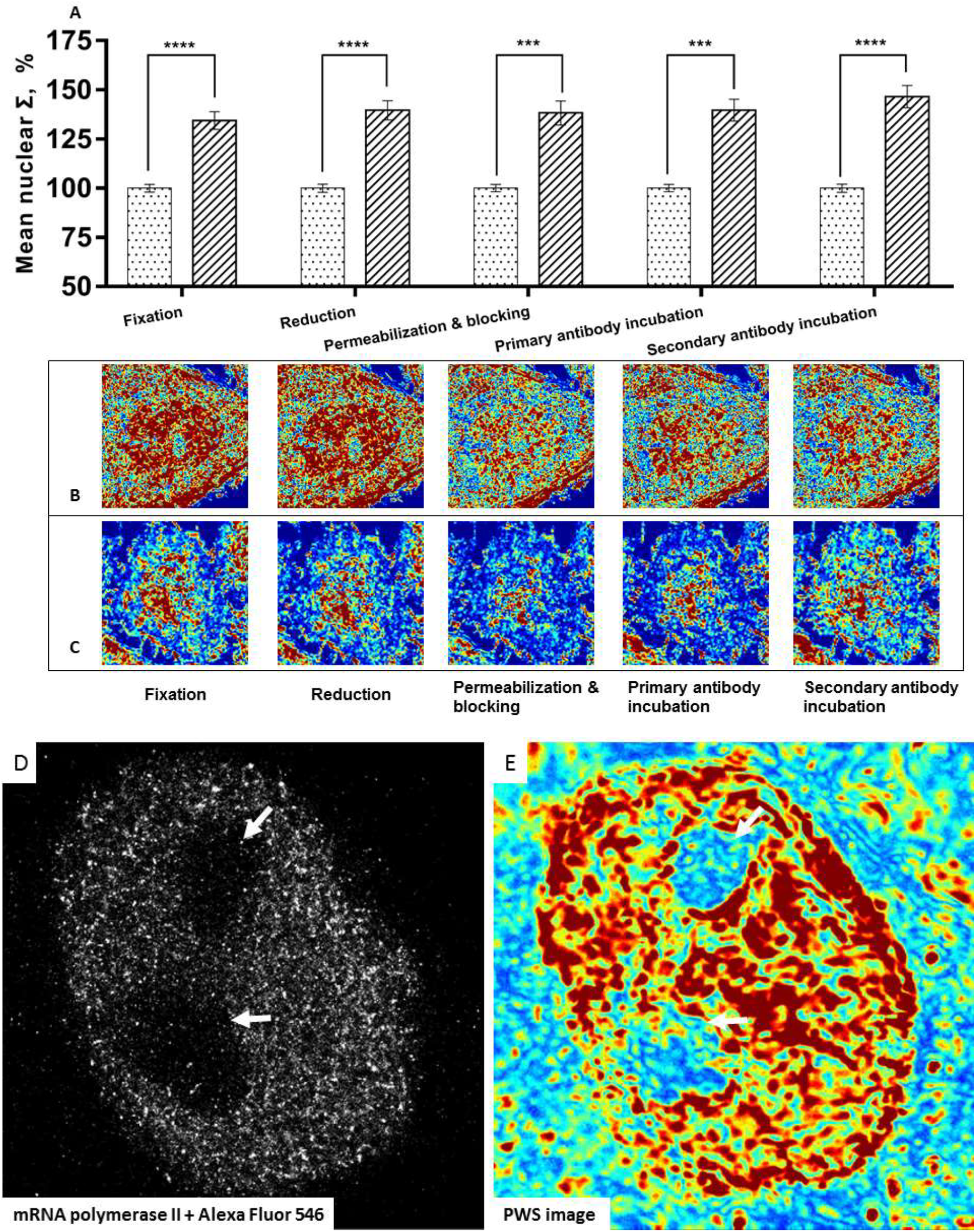
Effects of STORM preparation on chromatin ultrastructure. Human ovarian cancer cells were stained using antibody for mRNA polymerase II (Alexa Fluor 546). (A) Quantification of Σ (M248= 63 cells, A2780=71 cells) at each step of super resolution immunofluorescent labeling. Representative PWS images of M248 (B) and A2780 (C) at each step of fluorescent labeling. Representative STORM (D) and PWS image (E) for the ovarian cell nuclei for verification of the nanoscopic labeling process. Similar nuclear features can be seen on both PWS and STORM images. Arrows indicate nucleolus.

As can be seen in Fig. 7E, the relative nuclear Σ (*Ʃ_r_*) increases very rapidly with relative fluorescent intensity (*I_r_*) in low *I_r_* regions, and plateaus at around 50% *I_r_*.

## Discussion

We have developed two cell line models with nuclear structural differences to study the effects of chemical fixation and staining for nanopathological preparation using PWS microscopy for quantitative evaluation of the underlying chromatin ultrastructure. The nuclear structural differences were introduced by two different methods: chemotherapy treatment in the Hela model and p53 mutation in the human ovarian cancer model. These structural differences were quantified in live cells as ΔΣ, which can directly probe supra-nucleosomal chromatin structure and directly correlates with global patterns in gene expression [18, 25]. The two models have demonstrated stability and reproducibility, and thus enabling the systematic study of nanoscale diagnostic information changes during fixation processes both for label-free optical methods even when paired with cytological staining (Fig.6) or immunofluorescence (Fig. 7). Likewise, these methods are suitable for super-resolution based microscopy for multiple chromatin markers (Fig 8). Notably, we show by direct analysis of the same cells before and after fixation, ethanol and aldehyde fixation are both able to preserve ΔΣ while resulting in minimal changes to the overall cellular morphology. Although it has been shown that chemical fixatives would result in both cellular composition and structural changes [26, 27], our results indicate that the relative structural information at the nanoscale could be preserved during fixation. Therefore, the detection of nanoscale changes would still be possible despite the structural transformation. In studies of the subsequent preparation procedures after ethanol fixation, we found that a gradual rehydration and controlled air drying were both necessary to maintain differences in chromatin organization. Additionally, the application of histological staining and immunofluorescence can be used to demarcate the nucleus even though differences in ultra-structure are reduced between samples. This information loss could be attribute to the hydration and dehydration steps during staining which could alter the cellular structure at the nanoscale.

Besides ethanol-based fixation process, we also studied the immunofluorescent labeling process after aldehyde fixation. Immunofluorescence is a powerful tool in detecting the existence and spatial distribution of antigens. It is widely used in histopathology and is the basis for many super-resolution imaging techniques. However, the immunofluorescent labeling process involves multiple steps, which until now have not been validated rigorously between live and fixed cells at the nanoscale. It is known that these steps could largely determine the staining outcomes, but their effect on nanoscale diagnostic information was unstudied. In order to validate this process, we quantified changes in chromatin scaling of the same cells during all stages of the labeling process, and performed fluorescent microcopy and STORM at the end of the preparation. Our results show that the structural differences can be preserved during the labeling process for super-resolution imaging (Fig. 8). Finally, as the underlying structure of chromatin will likely vary based on molecular alterations, we demonstrate the capacity for direct co-localization analysis of chromatin domains. Notably, we found a clear trend between Σ and labeling density of H3K9me3 in the colocalization using PWS and fluorescent microscopy, which is frequently altered in carcinogenesis[28]. Consequently, these findings indicate that high throughput molecular-structural analysis could be useful for future nanopathological applications. In sum, this work enables the future detection of nanoscale structural alterations in human disease using nanocytological preparatory methods that are verified to represent those underlying a pathological process found in live cells.

## Conclusion

The emergence of imaging and molecular techniques capable of measuring the nanoscale structure of cells has the potential to greatly expand our understanding of biological function and human diseases. In the evaluation of human tissue, there are numerous preparatory steps required during the collection and processing of cells for nanoscopic analysis. Indeed, while nanopathology will likely emerge as a major advance in the near future, these capabilities depend on validated methods that can reliably maintain nanoscopic information between live and prepared cells. In this work, we demonstrate that although fixation, immunofluorescence, and histo-pathology staining alter structural information at the nanoscale, these methods can still reliably provide nanoscopic information for multiple imaging techniques. In particular, we use a recently developed imaging technique, live-cell PWS microscopy, to systematically study the effects of fixation and other preparatory processes on nuclear structure at the nanoscale. By using two distinct cell line models, we show that the relative structural information (ΔΣ) was well preserved during ethanol and aldehyde fixation, and during the subsequent preparatory steps required for super resolution fluorescent nanoscopy, immunofluorescent microscopy, and conventional histological examination. These findings would also be crucial for other fixed-cell based technologies with nanoscale sensitivity. In total, we developed a robust nano-cytological preparation process that can preserve the chromatin ultrastructure during all preparatory stages, from fixation to air drying, that can be applied as a standard validation for nanoscopic studies. This approach has proven effective in long term preservation of chromatin organization differences and it can be potentially applied in clinical nanoscopic studies. As this field will likely emerge due to the rapid pace of nanoscopic imaging techniques, this work provides a method to verify that studies of pathological processes extend to the structural behavior of living cells.

## Acknowledgement

We thank Di Zhang and Scott Gladstein for support and critical discussions. This material was based on work supported by National Institutes of Health (NIH)Grants R01CA155284, R01CA200064 and R01CA165309, and Lungevity Foundation.

---

**Fig S2.**
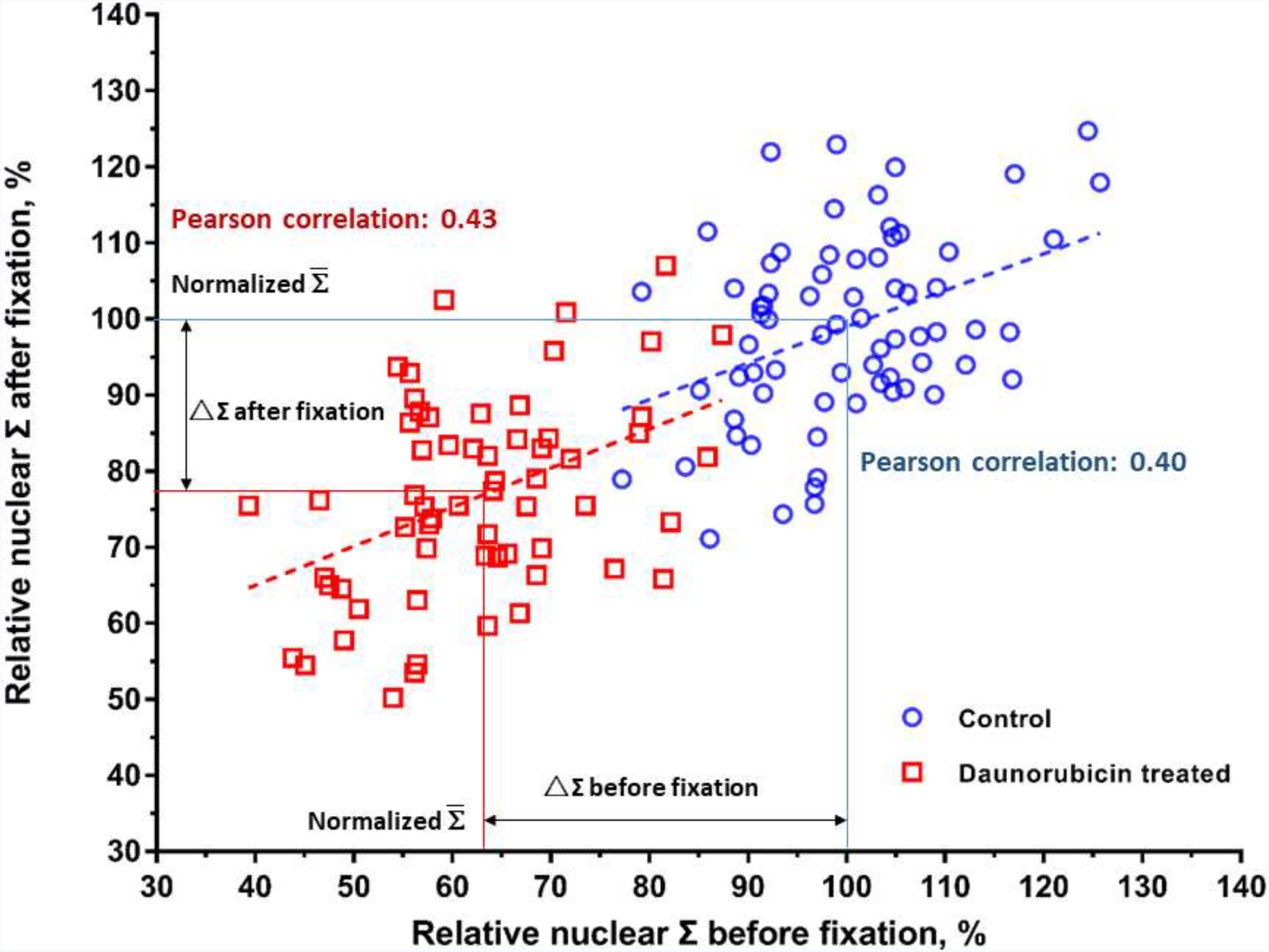
Correlation between relative nuclear Σ during 95% ethanol fixation. The same cells were tracked and imaged before and after 95% ethanol fixation (HeLa-control=71 cells, HeLa-daunorubicin=61 cells). The relative nuclear Σ was weakly correlated, but the population difference was preserved.

**Figure S5.**
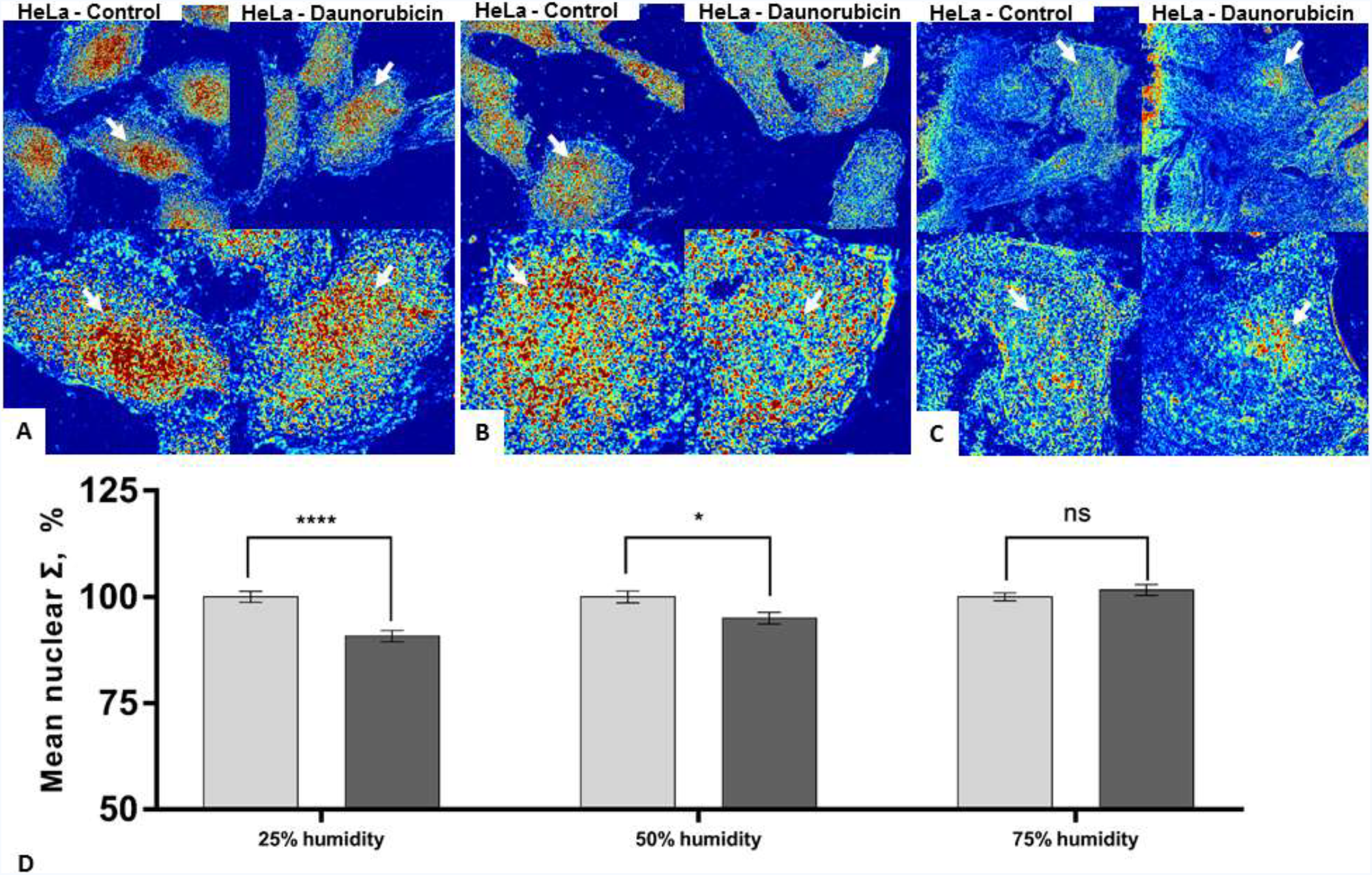
The effects of direct air drying on ΔΣ in HeLa model under varied humidity. Representative images of HeLa model air dried in (A) 25% (±5%), (B) 50% (±5%) and (C) 75% (±5%) humidity. (D) Quantification of mean nuclear Σ (25% humidity: HeLa-control=67 cells, HeLa-daunorubicin=72 cells; 50% humidity: HeLa-control=131 cells, HeLa-daunorubicin=141 cells; 75% humidity: HeLa-control=135 cells, HeLa-daunorubicin=126 cells) with SE bars.

